# Microbiome differentiation between micro-sympatric maize and teosinte reveals domestication-driven functional erosion of the microbiome across plant compartments

**DOI:** 10.1101/2025.10.14.682420

**Authors:** Esaú De la Vega-Camarillo, Shravan Sharma-Parunandi, Cesar Hernández-Rodríguez, Sanjay Antony-Babu, Julio S. Bernal

**Author notes:** **Corresponding author and email address:** Julio S. Bernal Sanjay Antony Babu.

## Abstract

**Introduction:** Crop domestication has fundamentally transformed plant phenotypes through artificial selection, yet the consequences of domestication for plant-associated microbial communities across the plant-soil continuum remain poorly understood.

**Gap Statement:** While recent studies suggest that domestication impacts microbiome structures, the magnitude, mechanisms, and functional implications of such impacts have not been systematically quantified using controlled experimental designs that eliminate environmental confounding factors.

**Aim:** To characterize and quantify the effects of crop domestication on microbial community structure and function by comparing maize (*Zea mays* subsp. *mays*) and its wild ancestor Balsas teosinte (*Zea mays* subsp. *parviglumis*) across multiple plant compartments in an unmanipulated field setting in Mexico, maize’s domestication center.

**Methodology:** We applied a micro-sympatric design in a natural setting to compare microbial communities between maize and Balsas teosinte across five plant compartments: bulk soil, rhizosphere, mucilage, leaves, and seeds. Full-length 16S rRNA gene sequencing was used for taxonomic characterization, Functional Annotation of Prokaryotic Taxa (FAPROTAX) and PICRUSt 2.0 were used to predict functional profiles, and network analysis was used to assess functional connectivity.

**Results:** Compartment identity explained 72.2% of variation in community structure, with consistent host effects across all niches (9.0%). Teosinte maintained significantly higher microbial diversity than maize across all compartments, with pronounced differences in seeds (32.0 ± 1.9 vs 9.3 ± 1.8 species, P < 0.01) and rhizosphere (60.3 ± 5.8 vs 33.8 ± 10.4 species, P < 0.01). Eighty-nine percent of predicted metabolic functions showed significant changes associated with domestication, with teosinte exhibiting enhanced nitrogen fixation (0.89 ± 0.07 vs 0.44 ± 0.04 in maize mucilage), siderophore production, and pathogen suppression. Network analysis revealed functional fragmentation in maize, with reduced connections (80 to 49) and lower clustering coefficients (0.62 ± 0.03 vs 0.25 ± 0.02, P < 0.001).

**Conclusion:** Balsas teosinte domestication fundamentally eroded microbial diversity and functional capacity in maize leading to a “domestication gap” that encompasses taxonomic loss, functional simplification, and network fragmentation, and replaced mutualistic plant-microbe partnerships with simplified microbial assemblages that may compromise crop resilience vis-à-vis a changing climate.

**Impact statement:** Understanding how plants select their microbial partners is crucial for enhancing agricultural productivity yet distinguishing between environmental and host genetic effects on microbiome assemblage remains challenging. Our study provides compelling evidence for host-driven microbiome assembly by comparing ancestral Balsas teosinte with derived maize growing in a common farm field in Mexico, eliminating environmental variation and experimental manipulation as confounding factors. By characterizing bacterial communities across different plant compartments, from soil to seed, we showed that each host’s genotype shaped divergent microbiome compositions despite growing in common environmental conditions. This research represents a significant step forward in understanding plant-microbe co-evolution during crop domestication and has three key implications. First, it suggests that microbiome traits were likely selected in conjunction with plant (host) traits during domestication and post-domestication selection. Second, it identifies specific bacterial communities that could be targeted for improving crop productivity and resilience. And third, it provides a methodological framework for studying host-microbe interactions in other crop-wild ancestor pairs. Our findings are particularly relevant for developing microbiome-based agricultural technologies and conservation strategies for beneficial plant-microbe interactions for deployment in traditional and modern farming systems.

**Data summary:** **The authors confirm all supporting data, code and protocols have been provided within the article or through supplementary data files.**

## 2. Introduction

Plant microbiomes fundamentally influence host survival and reproduction through multiple mechanisms, including nutrient acquisition, phytohormone modulation, immune system priming, and abiotic stress tolerance, among others [1–3]. While environmental factors strongly shape microbial communities [4,5], mounting evidence suggests that host genetic factors play a crucial yet poorly understood role in microbiome assembly [6,7]. An apparent interplay between host genetics and microbial recruitment has emerged as a key factor in plant evolution and adaptation and has significant implications for enhancing agricultural productivity in the face of environmental change [8,9, 125].

The domestication of Balsas teosinte (*Zea mays* ssp. *parviglumis*) leading to maize (*Zea mays* ssp. *mays*) is one of the best studied examples of crop domestication [10,11,126]. The morphological and genetic divergence between the two, beginning as early as 12,000 years ago in Mexico’s Balsas River Basin [10], was accompanied by significant changes in root architecture [13], exudate composition [14], and defensive compounds [15,16], among other traits. These modifications likely influenced plant-microbe interactions as recent studies demonstrated domestication-associated changes in defense-related metabolites [17] and root exudation patterns [18,19], among others.

The volume of research exploring domestication effects on plant-microbiome relationships has expanded significantly in recent years. Comparative studies in wheat [20], rice [21], and common bean [22] have demonstrated distinct microbial communities between crops and their wild relatives. In maize specifically, several studies have documented differences in rhizosphere communities between modern varieties and teosinte [23,24,127], though environmental variables, such as different growing conditions, confound most comparisons [25].

Plant-microbe interactions occur across distinct plant compartments, each compartment representing a unique ecological niche with selection pressures [26,27]. The rhizosphere hosts diverse microbial communities shaped by the soil and plant root exudates and immune responses [28,29]. Mucilage, a specialized polysaccharide-rich layer coating aerial roots of maize and other plants, represents an understudied compartment potentially crucial for microbial recruitment [30,31]. This mucilaginous layer shows structural and compositional differences between maize and teosinte [32], suggesting differential microbial recruitment. The phyllosphere, comprising above-ground plant surfaces, harbors distinct bacterial communities that influence plant health through various mechanisms [33,34].

Recent technological advances have accelerated our understanding of plant-microbe interactions. Full-length 16S rRNA sequencing provides improved taxonomic resolution compared to short-read approaches [35,36], enabling more precise characterization of bacterial communities. These methodological improvements, coupled with advanced bioinformatics tools [37,38], allow better prediction of functional capabilities from taxonomic data [39], though such predictions require validation through complementary approaches [40,41].

Studies comparing maize and teosinte microbiomes have typically focused on single compartments or plants grown in non-native environments. Several researchers have documented seed endophyte conservation across *Zea* species [42,43], while others have explored rhizosphere community differences between modern maize varieties and teosinte [44,45]. Investigations into root-associated microbiomes showed shifts in bacterial diversity associated with domestication [46,47], suggesting possible human-mediated selection on plant-microbe interactions. However, studies examining multiple plant compartments under identical environmental conditions are presently unavailable [48,49].

The traditional focus on rhizosphere communities has left significant knowledge gaps regarding other plant compartments [50,51]. The mucilage secreted by aerial roots represents an understudied interface that may play a crucial role in microbial recruitment and establishment [52,53]. Additionally, the relationship between phyllosphere communities and plant genotypes warrants a better understanding [54,55], especially in the context of crop domestication and subsequent selection and breeding [56].

In this study, we used a micro-sympatric field design to compare microbial communities between maize and Balsas teosinte across five plant compartments, viz. bulk soil, rhizosphere, mucilage, leaves, and seeds, in their native habitat in Mexico. Using full-length 16S rRNA gene sequencing and functional prediction approaches, we aimed to: (1) quantify host-driven effects on microbiome assembly across the plant-soil continuum, (2) identify core and host-specific bacterial communities, (3) assess functional consequences of domestication on plant-microbiomes interactions, and (4) evaluate network connectivity changes in microbial functional networks.

## 3. Methods

### Sample Collection and Processing

Field sampling was conducted in the community El Cuyotomate (19°58′14′′N, 104°04′ 06′′′W), Jalisco state, Mexico, in mid-September and mid-November 2024. For each of two plant types, Tuxpeño landrace maize and Balsas teosinte. We collected samples from four distinct compartments in mid-September: bulk soil (collected 30 cm from the plant), rhizosphere (soil firmly attached to roots), mucilage (gel-like substance coating the roots), and leaves (third fully expanded leaf) [57]. Subsequently, in mid-November we collected seed, a fifth compartment. Four biological replicates were collected per plant type and compartment. Samples were immediately preserved in DNA/RNA Shield™ (Zymo Research, USA) at a 1:3 ratio and transported to the laboratory at 4 °C [58].

### DNA Extraction and Amplification

Total genomic DNA was extracted from 0.5 g of solid samples (soil, leaves, seeds) or 0.5 mL of liquid samples (mucilage) using the ZymoBIOMICS™ DNA Miniprep Kit (Zymo Research, USA) according to the manufacturer’s instructions [59]. DNA quality and quantity were assessed using NanoDrop™ and Qubit™ fluorometric quantification [60].

The full-length 16S rRNA gene was amplified using universal bacterial primers 27F (5’-AGAGTTTGATCMTGGCTCAG-3’) and 1492R (5’-TACGGYTACCTTGTTACGACTT-3’) [61]. PCR reactions were performed in triplicate 25 μL volumes containing 12.5 μL Q5 Hot Start High-Fidelity Master Mix, 1.25 μL of each primer (10 μM), 2 μL template DNA, and nuclease-free water to volume [62]. Amplification success was verified using gel electrophoresis with 1% agarose gels.

### Library Preparation and Sequencing

PCR products were purified using AMPure XP beads (Beckman Coulter) with a 0.8X ratio [63]. Purified amplicons were quantified and standardized to 200 ng of DNA. Libraries were prepared using the Rapid Barcoding Kit (SQK-RBK114) following Oxford Nanopore’s specifications [64]. Sequencing was performed on a MinION Mk1B device using R9.4.1 flow cells for 72 hours [65].

### Bioinformatic Processing

Raw fast5 files were called using Guppy v6.0.1 (Oxford Nanopore Technologies) with the high-accuracy model [66]. Adapters and barcodes were removed using Porechop v0.2.4 with default parameters [67]. Quality filtering was performed using Prinseq++ v1.2, removing reads shorter than 1300 bp or longer than 1700 bp and sequences with mean quality scores below Q10 [68].

### Taxonomic and Diversity Analysis

Processed sequences were analyzed using the EZBioCloud platform [69]. Sequences were clustered into Operational Taxonomic Units (OTUs) at 99% similarity threshold for species-level resolution. Taxonomy was assigned using the EZBioCloud 16S database [70]. Alpha diversity metrics (Shannon, Chao1, and Simpson indices) were calculated after rarefying to even depth across samples. Beta diversity was assessed using weighted and unweighted UniFrac distances [71].

### Core and Specific Bacteriome Determination

Core bacteriome was defined as species (99% similarity threshold) present in 100% of samples within each compartment with significant relative abundance [72]. For specific bacteriome identification, we employed a dual approach: Linear discriminant analysis Effect Size (LEfSe) with LDA score > 2.0 and p < 0.05 [73] and XOR analysis to identify mutually exclusive species between plant types and compartments [74].

### Functional Predictions

Metabolic capabilities were predicted using two complementary approaches: FAPROTAX [75] for broad functional group assignment and PICRUSt2 [76] for more detailed metabolic pathway prediction. FAPROTAX classifications were based on curated taxonomic-functional associations, while PICRUSt2 predictions used the default KEGG Orthology database. Relative abundances were log-transformed and aggregated by compartment and host (maize vs. teosinte). A two-way ANOVA followed by Benjamini–Hochberg correction was applied to determine significant differences in functional enrichment across compartments. Functional traits with a false discovery rate (FDR) < 0.05 were retained for further interpretation [76]. To evaluate host-driven microbial recruitment, recruitment efficiency was defined as the ratio of rhizosphere to bulk soil diversity (Shannon Index). This metric was computed for each host and compared using Welch’s t-tests. Functional profiles were also analyzed using a log-ratio approach (e.g., nitrogen fixation traits vs. total predicted pathways), and were compared across host genotypes using bootstrapped confidence intervals (n = 999 iterations) [77]. Functional shifts were classified as “gained,” “conserved,” or “reduced” based on fold change and statistical significance (FDR < 0.05) between maize and teosinte for each function in each compartment. A ternary classification was used to quantify the proportion of functions within each category. Visual representation was performed using ggplot2 in R (v4.3.2), with significance indicated by asterisks [78].

### Network Construction and Topological Analysis

Microbial co-occurrence networks were constructed separately for each plant genotype (teosinte and maize) and compartment (root, rhizosphere, mucilage, and seed) using genus-level relative abundance tables. Prior to network inference, features with less than 0.01% mean relative abundance across samples or present in fewer than 20% of replicates were filtered to reduce noise [79].

Co-occurrence patterns were inferred using SparCC (Sparse Correlations for Compositional data) to account for compositionality in microbial community data. Correlations with absolute values ≥ 0.3 and adjusted *p*-values ≤ 0.05 (Benjamini-Hochberg FDR correction) were retained as significant edges. The resulting undirected networks were visualized and analyzed using the igraph and ggraph R packages. Key topological metrics—including node degree, edge density, clustering coefficient, modularity, and average path length—were calculated for each network to assess complexity and functional cohesion [80].

To identify compartment-specific microbial modules, we applied Louvain community detection. Ecological roles of taxa within modules were inferred based on existing literature and database annotations (KEGG, MetaCyc). Centrality measures (betweenness, closeness, and eigenvector centrality) were used to assess the keystone potential of individual genera [81].

### Statistical Analysis

All statistical analyses were performed using Python libraries (NumPy, SciPy, Pandas, and Scikit-bio) [82]. Data normality was assessed using the Shapiro-Wilk test, and homogeneity of variances was evaluated using Levene’s test [83]. Based on data distribution, appropriate statistical tests were selected: parametric (ANOVA with Tukey’s post-hoc) or non-parametric (Kruskal-Wallis with Dunn’s post-hoc) tests. Multiple testing corrections were performed using the Benjamini-Hochberg procedure [84].

## 4. Results

### Plant Compartment and Host Identity Drive Distinct Patterns of Microbial Alpha and Beta Diversity

We examined microbial diversity patterns across five plant compartments (bulk soil, rhizosphere, mucilage, leaves, seeds) in teosinte and maize using complementary alpha and beta diversity analyses (Figure 1). Alpha diversity metrics revealed significant compartment-specific differences, with species richness showing a clear hierarchical pattern across compartments (Figure 1a). Teosinte rhizosphere communities exhibited the highest richness (60.3 ± 5.8 species), with teosinte richness 1.8-fold greater than maize richness (p < 0.01). Bulk soil showed intermediate and similar richness values for both plants (teosinte: 42.0 ± 7.6; maize: 36.0 ± 3.8), while seeds and leaves displayed lower richness, with teosinte seeds maintaining 3.4-fold higher diversity than maize seeds (p < 0.01). Shannon diversity (Figure 1b) followed similar patterns, with teosinte consistently showing higher values across compartments, particularly in rhizosphere (H’ = 3.1 ± 0.3 vs. 2.6 ± 0.2, p < 0.05) and leaves (H’ = 1.5 ± 0.5 vs. 0.6 ± 0.2, p < 0.05).

**Figure 1.**
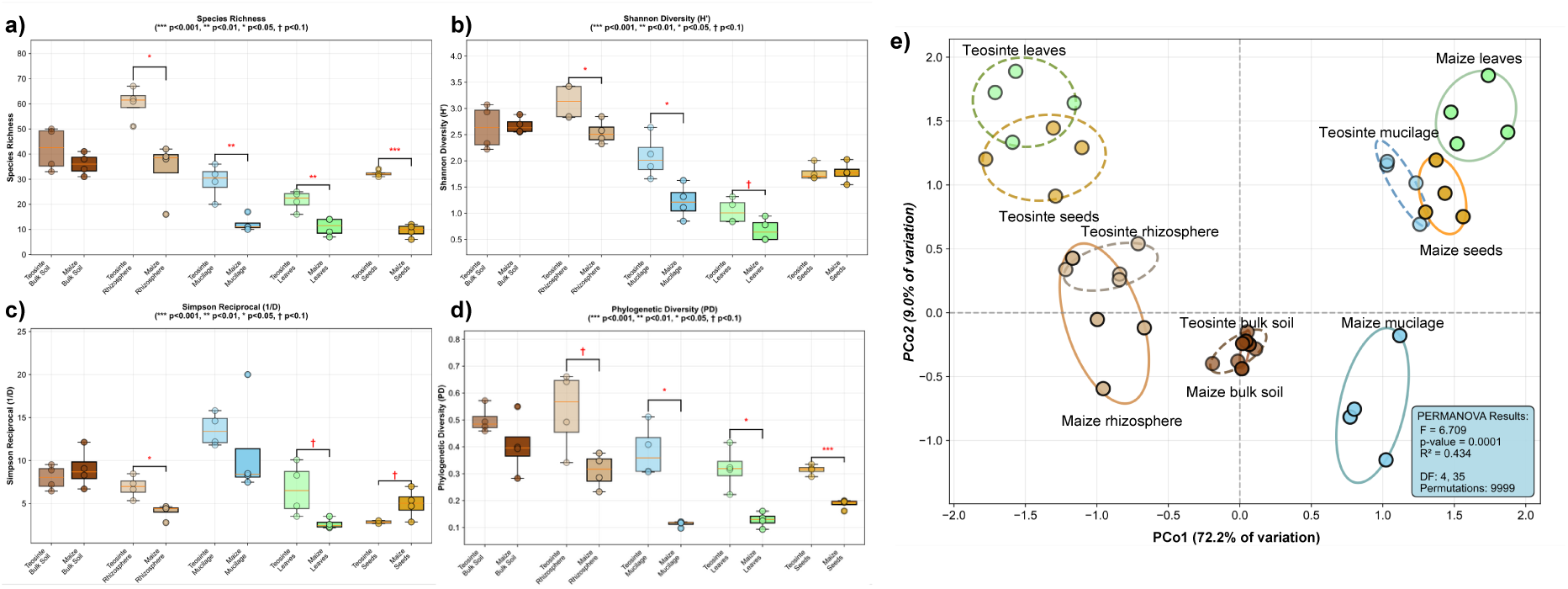
Alpha and beta diversity patterns reveal host-driven selection gradient across plant compartments. a-d) Alpha diversity metrics across five plant compartments in teosinte (light colors) and maize (dark colors): a) Species richness, b) Shannon diversity (H’), c) Simpson reciprocal diversity (1/D), and d) Phylogenetic diversity (PD). Boxplots show median, quartiles, and individual data points (n=4 biological replicates per group). Significance levels: ***p<0.001, **p<0.01, *p<0.05, †p<0.1. e) Principal coordinate analysis (PCoA) based on Bray-Curtis dissimilarity showing community structure across compartments. Ellipses represent 95% confidence intervals around group centroids. PERMANOVA results indicate significant effects of both compartment (F₄,₃₅ = 6.709, p < 0.0001, R² = 0.434) and host identity on microbial community composition. The analysis reveals a clear host-driven selection gradient from bulk soil (minimal host effect) through rhizosphere and mucilage (moderate host effects) to seeds and leaves (strong host effects), supporting the hypothesis that plant compartment specialization drives host-specific microbiome assembly.

Simpson reciprocal diversity (Figure 1c) and phylogenetic diversity (Figure 1d) revealed additional host-specific patterns, with mucilage showing pronounced differences between hosts (teosinte: 13.6 ± 1.7; maize: 11.1 ± 5.2). Chao1 estimator revealed distinct patterns across compartments, with teosinte showing higher richness in bulk soil (49.9 ± 5.1) and rhizosphere (66.0 ± 3.9) compared to maize (37.7 ± 4.1 and 39.9 ± 11.0, respectively, p < 0.05). This pattern reversed in seeds, where maize exhibited higher richness (38.3 ± 5.3) than teosinte (17.9 ± 2.0, p < 0.01) (Supplementary Figure 1a). Fisher’s alpha followed similar trends, with teosinte rhizosphere communities showing significantly higher estimated richness than maize (10.54 ± 0.8 vs. 6.5 ± 1.7, p < 0.01), while maize seeds again showed higher values (7.3 ± 1.0) compared to teosinte (2.4 ± 0.3, p < 0.001) (Supplementary Figure 1b). Pielou’s evenness showed relatively consistent values across most compartments for both hosts, ranging from approximately 0.2-0.4 in leaves to 0.8-1.0 in bulk soil and rhizosphere, with maize seeds showing notably higher evenness (0.8 ± 0.1) compared to teosinte seeds (0.3 ± 0.1, p < 0.001) (Supplementary Figure 1c).

Principal coordinates analysis (Figure 1e) revealed that host plant identity was the primary driver of community structure (PC1 = 72.2% of variation), horizontally separating teosinte leaf and seed samples (left) from corresponding maize samples (right), while compartment identity (PC2 = 9.0%) drove vertical separation of samples; maize and teosinte mucilage samples were separated vertically, unlike leaf and seed samples. The ordination displayed a compartment-driven vertical gradient, with bulk soil and rhizosphere samples clustering at the bottom, mucilage samples in the middle region, and seeds and leaves at the top. Within this structure, a host-driven selection gradient was evident, with bulk soil samples showing minimal separation between hosts, progressing through increasingly distinct host-specific clustering in rhizosphere and mucilage, culminating in seeds and leaves where teosinte and maize formed separate, well-defined clusters (Supplementary Figure 1d).

PERMANOVA confirmed significant effects of compartment (F₄,₃₅ = 6.709, p < 0.0001, R² = 0.434), host identity (p < 0.0001), and compartment × host interaction (p = 0.011) on community composition. The analysis revealed a clear host-driven selection gradient from bulk soil (non-significant host effect, p = 0.725) through rhizosphere (p = 0.012) and mucilage (p < 0.0001) to seeds and leaves (both p < 0.0001) (Supplementary Figure 1d). This pattern suggests that seeds and leaves represent particularly specialized communities with strong host specificity, potentially facilitating between-generation vertical transmission (i.e., from seed to seedling, or leaf litter to roots) of plant-adapted microorganisms and contributing to host-microbiome co-evolution during crop domestication.

### Core and Host-Specific Bacterial Communities

A compartment-resolved taxonomic analysis revealed distinct microbial assemblages in maize and teosinte, with clear differentiation between core and host-specific bacterial communities (Figure 2). Following normalization and quality control, we identified 13,085 total taxa across all compartments, with 1,038 taxa meeting significance criteria for categorization (7.9% of total diversity).

**Figure 2.**
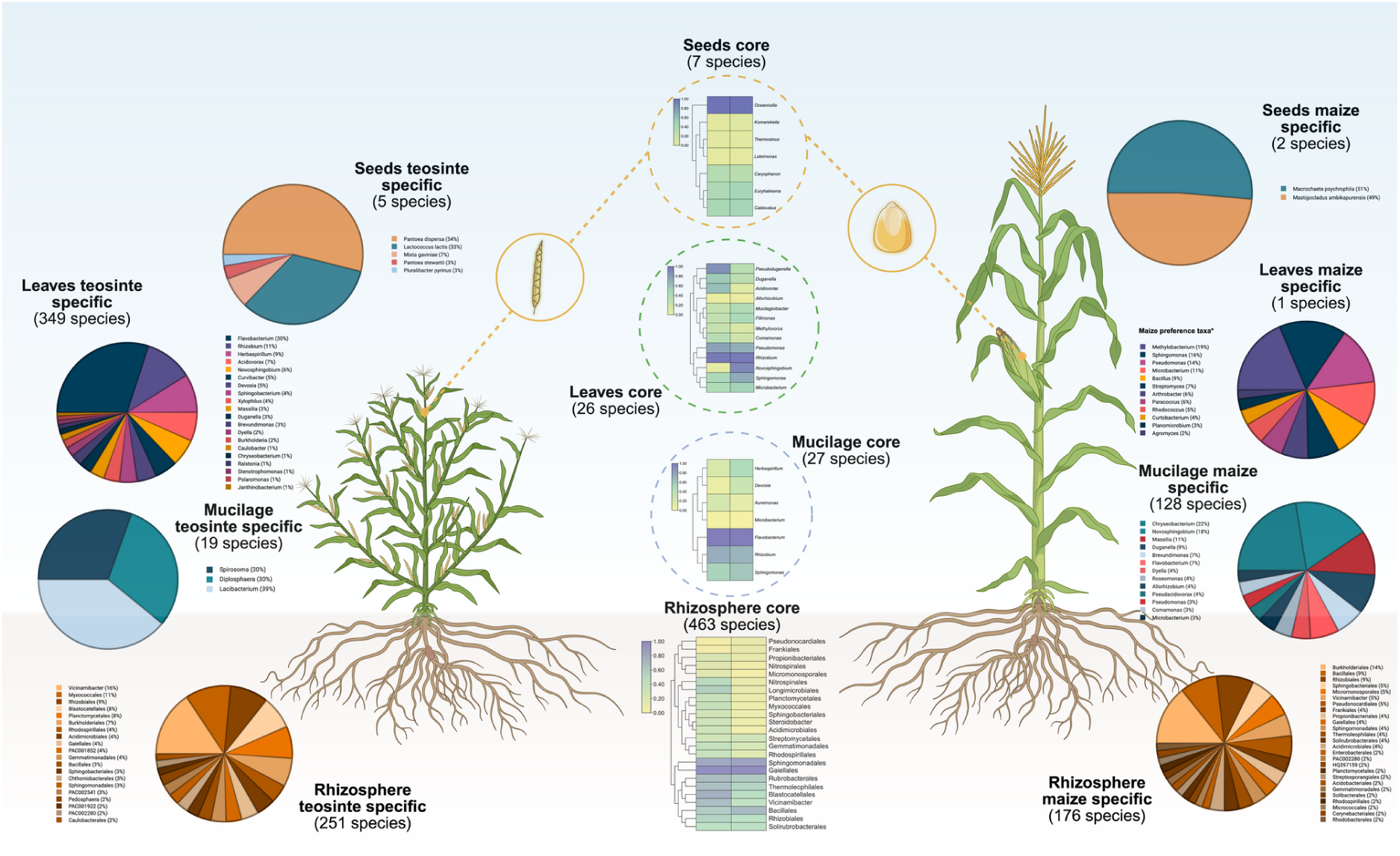
Compartment-specific bacterial community composition in maize and teosinte microbiomes. Taxonomic composition and abundance patterns across four plant-associated compartments (rhizosphere, mucilage, leaves, and seeds) showing core microbiome members shared between hosts and host-specific bacterial taxa. Pie charts display the relative abundance of dominant bacterial taxa in each compartment category: core communities (present in both hosts), teosinte-specific taxa (exclusive to teosinte), and maize-specific taxa (exclusive to maize). The central heatmaps show the taxonomic composition of core communities at the order level, with color intensity representing relative abundance. Plant illustrations indicate the anatomical compartments sampled: rhizosphere (root-associated soil), mucilage (root exudates and mucilaginous secretions), leaves (phyllosphere), and seeds (endosphere). Numbers in parentheses indicate the total number of bacterial species or genera identified in each category. The analysis reveals distinct microbial signatures between maize and teosinte across all compartments, with the rhizosphere showing the highest microbial diversity (1,420 total taxa) and seeds displaying the most pronounced host-specificity with minimal core overlap (7 shared species).

Rhizosphere communities exhibited the highest microbial complexity and most balanced host interactions. The maize-teosinte rhizosphere core bacteriome comprised 463 shared species (7.4% of 6,232 total taxa), representing the largest absolute core across all compartments. This core was dominated by *Rubrobacter* (abundance ratio M/T = 1.16), *Gaiella* (ratio = 1.06), and members of Sphingomonadales and Rhizobiales orders, reflecting conserved adaptation to root exudate environments. Host-specific differentiation was moderate but significant, with 176 maize-exclusive species (2.8%) and 251 teosinte-exclusive species (4.0%). Additionally, 77 maize-enriched (1.2%) and 71 teosinte-enriched (1.1%) species demonstrated intermediate host preferences, indicating gradual rather than binary host selection in the rhizosphere.

Mucilage communities showed the strongest maize-specific enrichment patterns. The combined maize-teosinte mucilage core bacteriome was restricted to 27 species (0.9% of 2,985 total taxa) but included functionally important genera: *Flavobacterium* (ratio M/T = 1.50), *Rhizobium* (ratio = 1.02), and *Herbaspirillum* (ratio = 0.84), suggesting conserved functions in root exudate processing and nitrogen cycling. Notably, maize mucilage harbored 128 exclusive species (4.3%) compared to only 19 teosinte-exclusive species (0.6%), representing a 6.7-fold bias toward maize specialization. This pattern was complemented by 8 maize-enriched (0.3%) and 19 teosinte-enriched (0.6%) species.

Leaf-associated communities displayed the most pronounced teosinte bias, with a combined minimal core of 26 species (0.7% of 3,849 total taxa) dominated by *Flavobacterium* (ratio M/T = 0.85), *Rhizobium* (ratio = 1.02), and *Novosphingobium* (ratio = 0.70). The striking asymmetry in host specificity was evident with 349 teosinte-exclusive species (9.1%) versus only 1 maize-exclusive species (0.03%), representing a 349-fold bias toward teosinte. This was accompanied by 12 maize-enriched (0.31%) and 2 teosinte-enriched (0.05%) species, suggesting that domestication has led to substantial simplification of leaf endosphere microbial communities in maize.

Seed-associated communities exhibited the highest proportional core diversity despite lowest absolute diversity. With 19 total taxa, seeds maintained a substantial, combined core of 7 shared species (36.8% of seed diversity), representing the highest core percentage across all compartments. This seed core was dominated by *Oceanicella actignis* (38.7% ± 4.1% relative abundance in maize vs 14.0% ± 2.2% in teosinte), *Caldovatus sediminis* (17.7% ± 2.3% in maize vs 6.4% ± 1.1% in teosinte), *Euryhalinema mangrovii* (17.3% ± 2.1% in maize vs 6.0% ± 0.9% in teosinte), and *Caryophanon latum* (12.4% ± 0.1% in maize vs 5.3% ± 0.4% in teosinte). Despite this substantial shared core, seeds also showed pronounced host-specific patterns with 5 teosinte-exclusive species (26.3%), including *Pantoea dispersa* (33.4% ± 1.1%), *Lactococcus lactis* (20.6% ± 0.7%), *Mixta gaviniae* (4.4% ± 0.3%), *Pantoea stewartii* (1.9% ± 0.3%), and *Pluralibacter pyrinus* (1.7% ± 0.2%). Maize seeds displayed 4 enriched species (21.1%), suggesting that while vertical transmission maintains a conserved core microbiome, host-specific selection has also driven divergent seed microbiome assembly.

### Domestication drives compartment-specific microbiome restructuring with functional consequences

We performed functional profiling to predict metabolic capabilities of bacterial communities across plant compartments in teosinte and maize (Figure 3). The analysis revealed distinct functional profiles between hosts and across compartments, with clustering patterns reflecting both phylogenetic relationships and environmental selection pressures.

**Figure 3.**
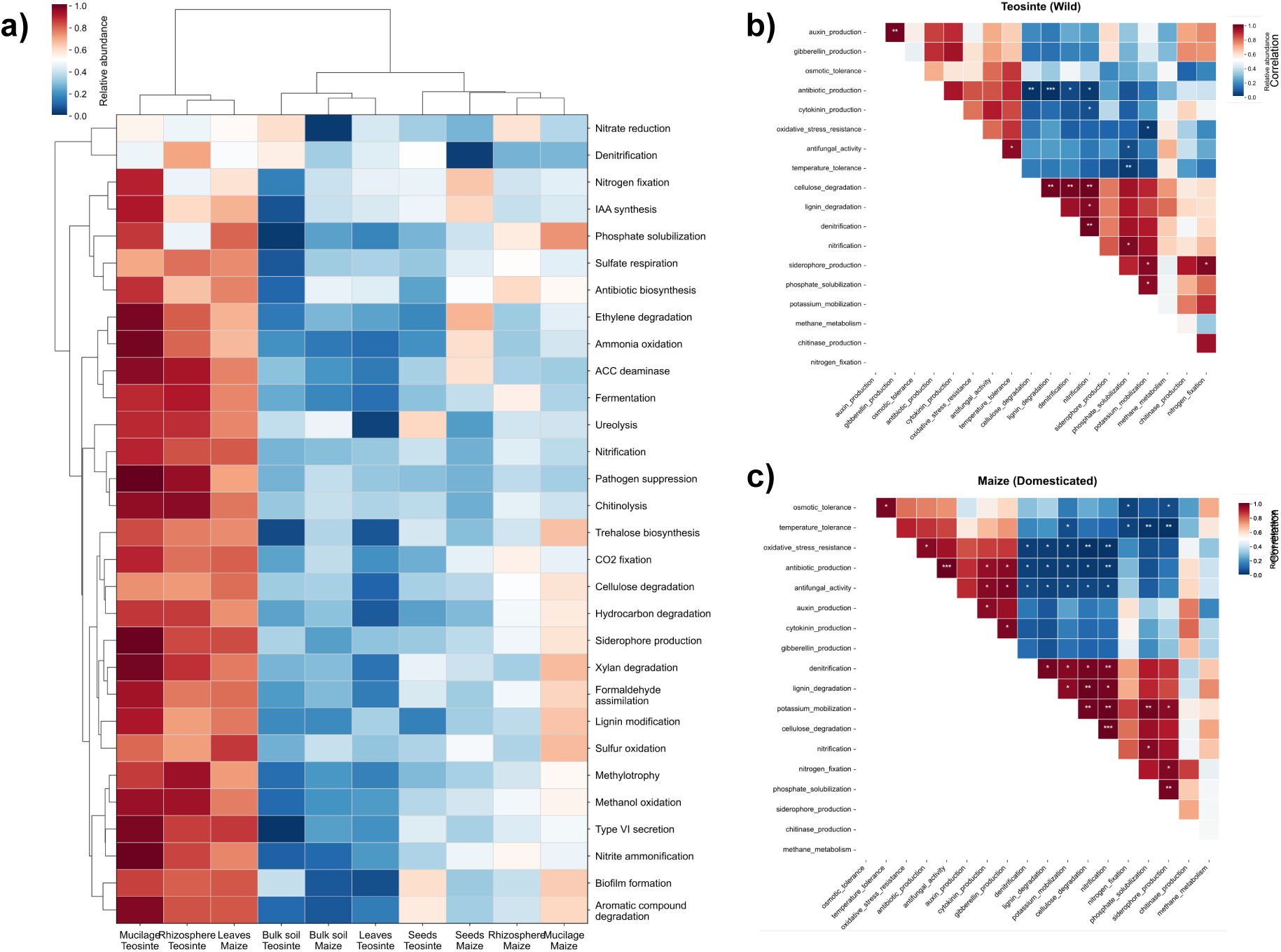
Functional profiling reveals compartment-specific metabolic capabilities and host-dependent functional specialization. a) Hierarchical clustering heatmap of predicted metabolic functions across plant compartments in both teosinte and maize, showing relative abundance of functional categories normalized across all samples. Clustering reveals compartment-driven functional groupings with distinct metabolic profiles. b-c) Correlation matrices between predicted metabolic functions and plant compartments for teosinte (b) and maize (c). Color intensity represents correlation strength (red = positive correlation, blue = negative correlation), with asterisks indicating statistical significance levels (*p < 0.05, **p < 0.01, ***p < 0.001). Functions include nitrogen cycling (denitrification, nitrogen fixation, nitrate reduction, ureolysis, nitrification), plant growth promotion (IAA synthesis, ACC deaminase, phosphate solubilization), plant defense (antibiotic biosynthesis, pathogen suppression, siderophore production), carbon metabolism (fermentation, cellulose degradation, xylan degradation), and stress responses (ethylene degradation, trehalose biosynthesis). Values represent mean ± standard deviation (n=4 biological replicates per group). The analysis reveals compartment-specific functional specialization with notable differences between teosinte and maize, particularly in mucilage communities where teosinte shows enhanced multifunctional capabilities, suggesting functional reorganization of microbial communities during crop domestication.

Hierarchical clustering of predicted metabolic functions (Figure 3a) shows the first cluster grouped teosinte mucilage and rhizosphere together with maize leaves, showing high relative abundances of multiple functional traits, with the strongest signals in teosinte compartments, all other compartments–bulk soil, seeds and maize rhizosphere and mucilage–formed a second, more heterogeneous cluster characterized by lower overall functional abundances.

Correlation analysis between predicted metabolic functions and plant compartments revealed several key patterns (Figure 3b, c). Teosinte mucilage communities showed the highest functional diversity, with peak activities in pathogen suppression (0.998 ± 0.050), siderophore production (0.992 ± 0.048), nitrite ammonification (0.991 ± 0.045), xylan degradation (0.980 ± 0.052), and ammonia oxidation (0.979 ± 0.048) (Figure 3b). In contrast, maize mucilage showed more moderate functional levels across most categories (Figure 3c). Rhizosphere communities in both hosts exhibited high nitrogen cycling capabilities, with teosinte showing particularly elevated levels in pathogen suppression (0.937 ± 0.065), chitinolysis (0.950 ± 0.058), and methylotrophy (0.926 ± 0.071), while maize rhizosphere displayed strong positive correlations with lignin degradation, denitrification, and potassium mobilization, alongside strong negative correlations with antibiotic and antifungal production.

Notable host-specific differences included enhanced nitrogen fixation in teosinte mucilage (0.890 ± 0.067 vs. 0.441 ± 0.044 in maize) and elevated ACC deaminase activity (0.952 ± 0.063 vs. 0.321 ± 0.042). Teosinte consistently showed higher denitrification potential across compartments, particularly in rhizosphere (0.696 ± 0.058 vs. 0.265 ± 0.035) and mucilage (0.462 ± 0.048 vs. 0.271 ± 0.038). Conversely, maize displayed enhanced antibiotic biosynthesis in leaves (0.749 ± 0.071 vs. 0.432 ± 0.055) and seeds (0.491 ± 0.058 vs. 0.232 ± 0.032), suggesting potential defensive adaptations with domestication.

Leaf communities showed distinct functional profiles between hosts, with maize leaves exhibiting higher sulfur oxidation (0.855 ± 0.078 vs. 0.303 ± 0.042), type VI secretion (0.853 ± 0.082 vs. 0.196 ± 0.028), and biofilm formation (0.812 ± 0.075 vs. 0.059 ± 0.015) compared to teosinte. Seed communities displayed relatively lower functional diversity overall, but maize seeds showed enhanced nitrogen fixation (0.638 ± 0.064 vs. 0.456 ± 0.048) and ethylene degradation (0.671 ± 0.063 vs. 0.177 ± 0.025) capabilities.

Bulk soil communities represented baseline functional profiles, with generally lower activity levels across most functions compared to plant compartments. However, teosinte bulk soil showed notably higher nitrate reduction (0.589 ± 0.058 vs. 0.020 ± 0.008) and denitrification (0.543 ± 0.054 vs. 0.326 ± 0.038) compared to maize bulk soil, suggesting host-driven effects extending beyond the rhizosphere and into the surrounding soil environment.

These functional patterns suggest that domestication altered the metabolic landscape of plant-associated microbial communities, with teosinte maintaining more diverse nitrogen cycling and plant growth-promoting functions, and maize evolving enhanced defensive and stress-response capabilities, potentially reflecting adaptation to agricultural environments and farmer-selection pressures.

### Domestication Drives Systematic Loss of Microbial Diversity and Functional Reorganization Across Plant Compartments

We analyzed phylogenetic diversity patterns and recruitment efficiency across plant compartments to assess any evolutionary implications of domestication on microbial community assembly (Figure 4). Between-plant comparisons of richness and phylogenetic diversity revealed a clear “domestication gap” across all plant compartments, with teosinte consistently maintaining higher richness and phylogenetic diversity than maize (Figure 4a).

**Figure 4.**
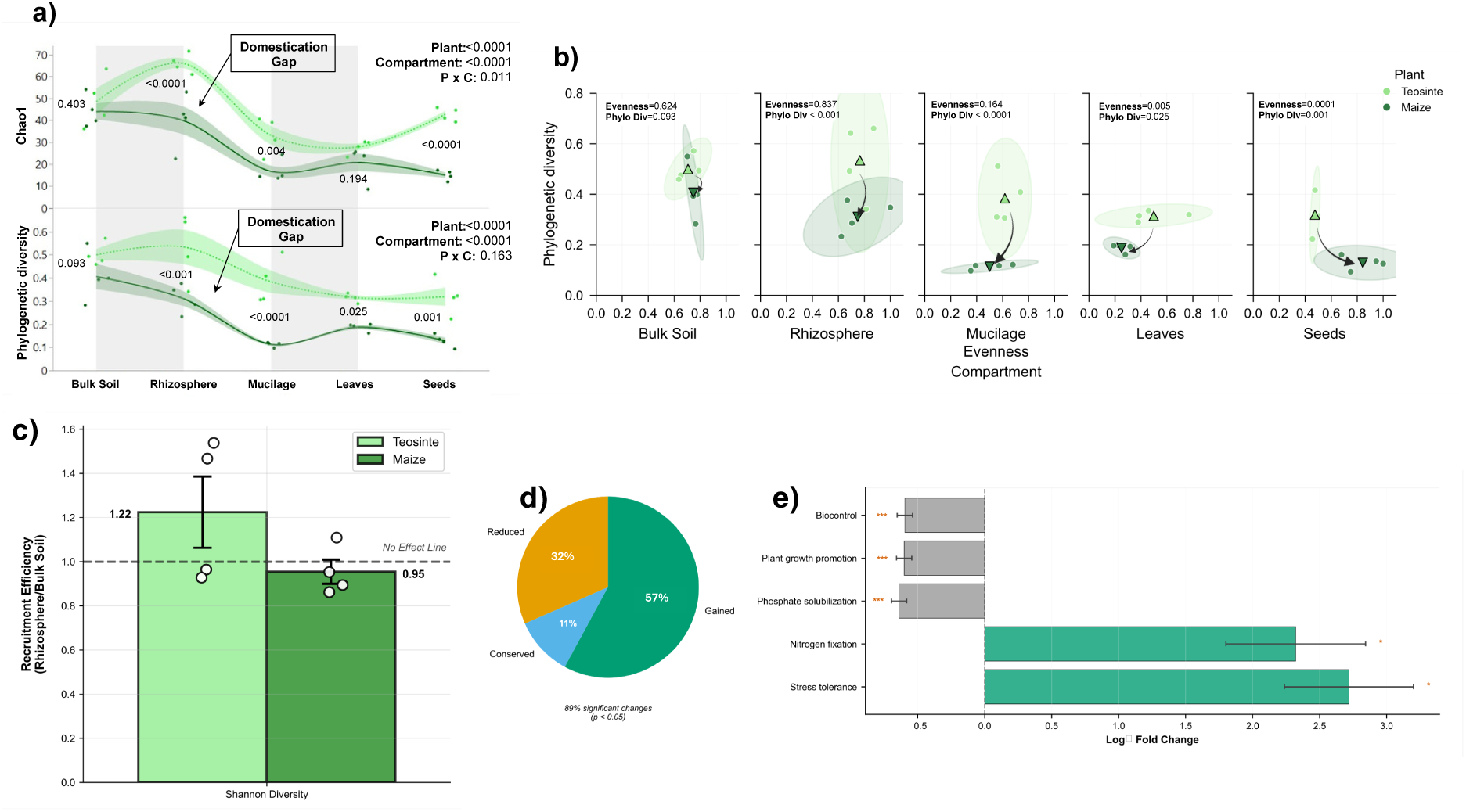
Domestication drives systematic loss of microbial diversity and functional reorganization across plant compartments. a) Phylogenetic diversity metrics (Chao1 and phylogenetic diversity) across plant compartments showing consistent domestication gaps between teosinte (light green) and maize (dark green). P-values indicate significance levels of differences between hosts within each compartment. Shaded areas represent 95% confidence intervals around regression lines. b) Shannon diversity vs. species richness scatter plot with 95% confidence ellipses showing domestication trajectory from teosinte (circles) to maize (squares). Colors represent different plant compartments (bulk soil, rhizosphere, mucilage, leaves, seeds). PERMANOVA results confirm significant plant and compartment effects with interaction. c) Recruitment efficiency comparing microbial community assembly capacity between teosinte and maize, calculated as the ratio of rhizosphere to bulk soil diversity (mean ± SE, n=4). Dashed line indicates no effect threshold (1.0). Individual data points show biological replicates. d) Pie chart summarizing functional changes with domestication based on FAPROTAX analysis, with 89% of predicted metabolic functions showing significant alterations (P < 0.05). e) Log fold change analysis of specific functional categories showing magnitude and direction of change from teosinte to maize. Enhanced functions in maize are shown in green (stress tolerance, nitrogen fixation), while reduced functions are shown in gray (biocontrol, plant growth promotion, phosphate solubilization). Error bars represent standard error; asterisks indicate statistical significance (*P < 0.05, **P < 0.01, ***P < 0.001). The analysis reveals that domestication has systematically reduced microbial diversity while reorganizing functional capabilities, with implications for crop resilience and sustainable agriculture.

**Figure 5.**
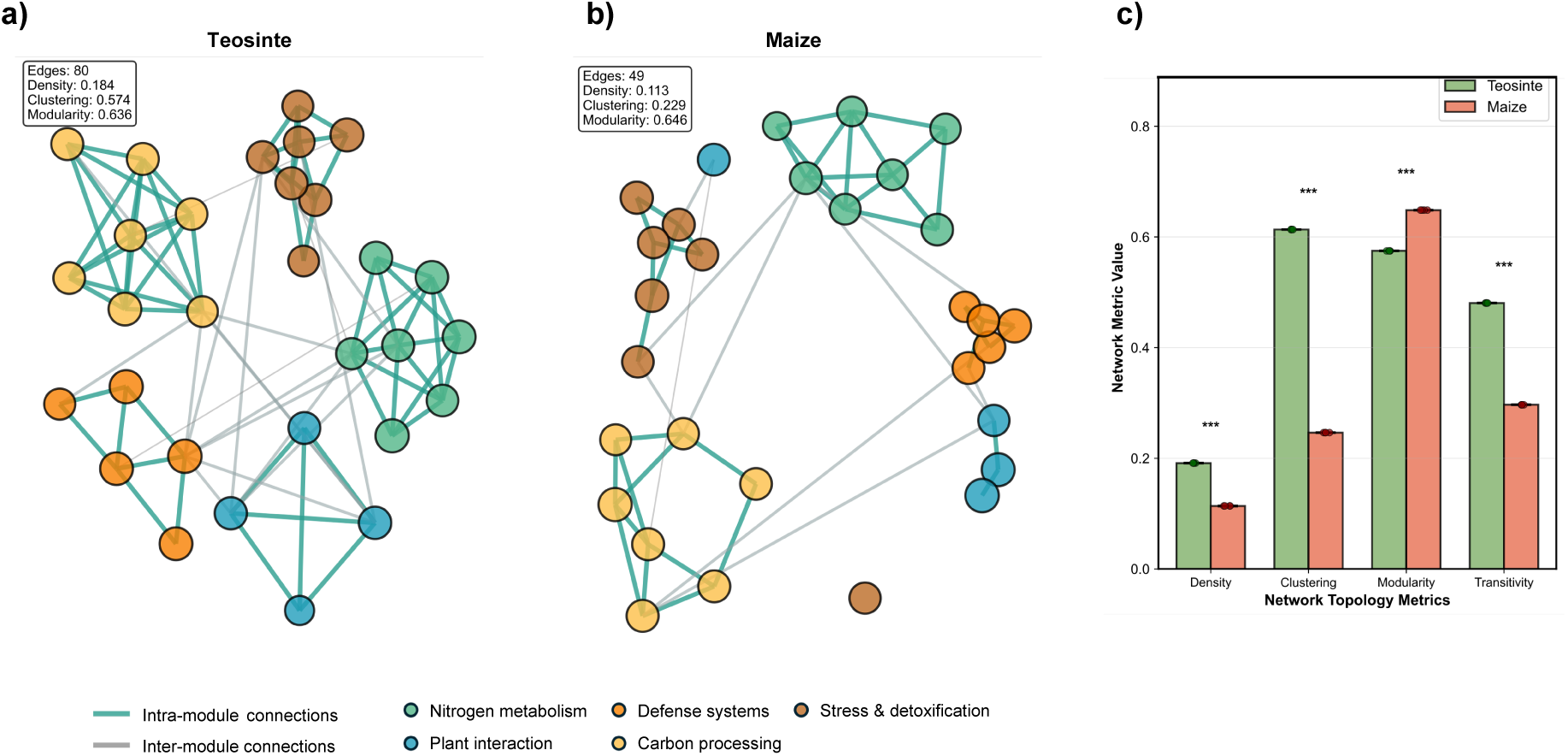
Functional Network Restructuring During Maize Domestication. Network analysis of 30 key microbial functions reveals dramatic restructuring of functional connectivity between teosinte (a) and maize (b) microbiomes. (a) Teosinte exhibits a highly modular network with 80 functional connections, strong clustering (0.574), and integrated functional modules represented by different node colors: nitrogen metabolism (green), plant interaction (blue), defense systems (orange), carbon processing (yellow), and stress & detoxification (brown). Thick teal edges represent intra-module connections, while gray edges show inter-module links. (b) Maize displays a fragmented network with only 49 connections, reduced clustering (0.229), and weakened inter-module connectivity, indicating loss of functional integration with domestication. Node sizes reflect relative functional activity across plant compartments. (c) Quantitative comparison of network topology metrics shows significant differences across all parameters (***P < 0.001, two-tailed t-test, n=4 biological replicates per host). Error bars represent standard error of the mean (SEM). Individual data points show replicate variation. The analysis demonstrates that domestication has led to substantial simplification of microbial functional networks, with reduced connectivity, clustering, and transitivity, potentially compromising the resilience and metabolic flexibility of agricultural microbiomes compared to their wild counterparts.

Across plant compartments, the domestication gap between teosinte and maize was significant and similar in size for both species richness and functional diversity (Figure 4a). Richness was ∼1.6-fold greater (P < 0.0001) and functional diversity ∼1.8-fold greater (P < 0.0001) in teosinte compared to maize. Between compartments and for species richness (Chao1), the domestication gap was greatest for the rhizosphere (F_1,30_ = 23.937, P < 0.0001) and leaf (F_1,30_ = 26.994, P < 0.0001) compartments, and not significant for bulk soil (P = 0.403) and leaves (P = 0.194). For functional diversity (phylogenetic diversity), the domestication gap was greatest for the mucilage compartment (F_1,30_ = 25.204, P < 0.0001), followed by rhizosphere, seed, and leaf compartments (P = 0.025 to < 0.001), and not significant for bulk soil (F_1,30_ = 3.020, P = 0.093). PERMANOVA analysis confirmed significant effects of both plant identity (P < 0.0001) and compartment identity (p < 0.0001), with a significant interaction effect (P × C: P = 0.011). These results reveal a clear domestication gap between teosinte and maize, i.e., a significant microbial diversity gap across plant-associated compartments between a crop and its wild ancestor.

Phylogenetic diversity plotted against evenness (Figure 4b) revealed distinct clustering patterns along a domestication trajectory. Teosinte samples occupied regions of higher phylogenetic diversity across most compartments, while maize samples clustered in areas of reduced phylogenetic diversity. The 95% confidence ellipses showed minimal overlap between hosts in mucilage (P = 0.011), leaves (P < 0.001), and seeds (P = 0.012), confirming significant divergence in phylogenetic diversity patterns. Domestication of teosinte seemed accompanied by a consistent reduction in phylogenetic diversity across plant-associated compartments during crop evolution, with bulk soil (P = 0.192) and rhizosphere (P = 0.050) showing less pronounced differences.

Recruitment efficiency analysis (Figure 4c) showed significantly higher efficiency in recruiting diverse microbial communities from the rhizosphere to bulk soil (1.22 ± 0.18) in teosinte compared to maize (0.95 ± 0.05). This difference suggests that teosinte retains superior capacity for assembling diverse microbial communities, potentially reflecting maintained co-evolutionary relationships that were disrupted with domestication in maize.

Functional change analysis revealed that 89% of predicted metabolic functions showed significant changes associated with domestication (p < 0.05). Of these changes in maize, 57% represented functional gains, 32% were reductions, and only 11% remained conserved between hosts (Figure 4d). This pattern suggests substantial functional reorganization of microbial communities in maize with domestication of teosinte.

Detailed functional analysis (Figure 4e) identified specific categories of change with domestication. Stress tolerance and nitrogen fixation functions showed the largest increases in maize (log fold change > 2.0, P < 0.05), suggesting enhanced adaptation to agricultural environments and potential compensatory mechanisms for reduced microbial diversity. Conversely, biocontrol, plant growth promotion, and phosphate solubilization functions were weaker in maize compared to teosinte, indicating potential loss of beneficial plant-microbe interactions. These functional shifts suggest that while some adaptive functions may have been enhanced with domestication, important plant growth-promoting and protective functions may have been compromised, potentially affecting plant fitness and resilience.

### Functional Network Restructuring During Maize Domestication

Network analysis of 30 key microbial functions revealed dramatic restructuring of functional connectivity between the teosinte and maize microbiomes. The teosinte-associated microbiome exhibited a highly modular functional network characterized by dense intra-module connectivity and strategic inter-module bridges, containing 80 functional connections with high clustering coefficient (0.574) and strong modularity (0.636). Five distinct functional modules were identified: nitrogen metabolism, plant interaction, defense systems, carbon processing, and stress and detoxification pathways. In contrast, the maize-associated microbiome displayed a significantly fragmented network architecture with 49 functional connections and substantially reduced clustering (0.229), indicating weakened functional integration between metabolic pathways.

Statistical comparison of network topology metrics revealed significant differences between teosinte and maize microbiomes across all measured parameters. Network density was significantly higher in teosinte (0.19 ± 0.01) compared to maize (0.13 ± 0.01, P < 0.001), while the clustering coefficient was dramatically reduced in maize, showing a 2.5-fold decrease from teosinte (0.62 ± 0.03) to maize (0.25 ± 0.02, P < 0.001). Network transitivity was also significantly higher in teosinte (0.49 ± 0.02) compared to maize (0.29 ± 0.02, P < 0.001), indicating weaker functional cooperation and pathway integration with domestication. These results demonstrate that domestication is associated with substantial simplification of microbial functional networks, with reduced connectivity, clustering, and transitivity, potentially compromising the resilience and metabolic flexibility of crop microbiomes compared to their wild counterparts.

## 5. Discussion

This study provides insight into how plant domestication systematically reconfigured plant-microbe relationships across the plant-soil continuum. By isolating host-genetic effects through a micro-sympatric field design and leveraging full-length 16S rRNA sequencing we revealed host compartment- and host-driven microbiome differentiation between maize and its wild ancestor Balsas teosinte. Our study quantified the magnitude and scope of microbial community restructuring associated with domestication and post-domestication selection by farmers, revealing a multidimensional erosion of microbiome complexity or “domestication gap”. These results not only demonstrate domestication effects at the microbiome level but also provide a quantitative framework for their understanding, with implications for crop resilience and agricultural sustainability in the face of climate change.

### Domestication and the Evolutionary Disruption of Microbial Assembly Rules

Our quantitative analysis reveals that host plant identity is the primary driver of variation in community structure, evident with its spread mostly along the horizontal axis explaining >70% of variation (PC1), while compartment was mostly spread along the vertical axis explaining <10% of variation (PC2) (see Figure 1e). This dominant host effect implies strong and statistically significant genetic control over microbial recruitment patterns across all plant niches. The host-driven selection gradient is particularly pronounced in specialized compartments such as seeds and leaves, where teosinte and maize form separate clusters (see Figure 1e), while being minimal in bulk soil where both hosts share similar microbial communities. This pattern suggests that plant-specific chemical traits (e.g., exudate profiles, mucilage composition) and vertical transmission mechanisms may play predominant roles in shaping host-specific microbial assemblages, particularly in above-ground tissues and seed-associated communities where host genetic effects are strongest (e.g., p < 0.0001 for both seeds and leaves; Supplementary Figure 1d). The systematic differences in microbial diversity between teosinte and maize across compartments— with teosinte maintaining higher richness in practically all niches suggests that teosinte domestication fundamentally restructured microbiome assembly patterns in maize rather than simply reducing overall diversity.

The loss of core mucilage taxa and their associated functions reflect a disruption of co-evolved diazotrophic interactions, potentially compromising associative nitrogen fixation [86–88]. This functional erosion is quantitatively supported by our findings showing teosinte mucilage communities maintaining significantly higher nitrogen fixation capacity (0.89 ± 0.07) compared to maize (0.44 ± 0.04), representing a 50% reduction in this critical plant growth-promoting function. Similarly, teosinte shows enhanced pathogen suppression (0.998 ± 0.050), siderophore production (0.992 ± 0.048), and ACC deaminase activity (0.952 ± 0.063 vs. 0.321 ± 0.042 in maize).

From an evolutionary standpoint, such shifts point to a decoupling of host-microbe coadaptation, where artificial selection—targeting yield, growth habit, and grain quality, among other traits—inadvertently neglected microbial recruitment traits, a phenomenon increasingly recognized across crops and their wild relatives [89]. This decoupling is most evident in plant-associated compartments, with the rhizosphere and seed microbiomes showing the most pronounced changes. Teosinte maintains significantly higher diversity in the rhizosphere (60.3 ± 5.8 species) compared to maize (33.8 ± 10.4 species, p < 0.01), and even more dramatically in seeds (teosinte: 32.0 ± 1.9 species vs. maize: 9.3 ± 1.8 species, p < 0.01). This pattern suggests that microbial inheritance pathways were bottlenecked during farmer selection that prioritized observable agronomic and quality traits over unseen microbial partnerships [128,129].

### The Domestication Gap: A Framework for Understanding Microbiome Erosion

We propose the term “domestication gap” to describe the across-compartments reduction in microbial diversity and functional capacity observed between crop species and their wild ancestors. This concept extends beyond simple diversity loss to encompass the multidimensional erosion of microbiome complexity during artificial selection [127–135]. Our analysis revealed that this domestication gap varied markedly across plant compartments, being most pronounced in rhizosphere (Chao1, p < 0.0001; phylogenetic diversity, p < 0.001) and seed (Chao1, p < 0.0001; phylogenetic diversity, p = 0.001) compartments, while remaining marginal in leaves (Chao1, p = 0.194; phylogenetic diversity, p = 0.025), and non-significant in bulk soil (Chao1, p = 0.403; phylogenetic diversity, p = 0.093), indicating that host-driven selection effects are strongest in plant-associated niches rather than in the surrounding soil environment.

The domestication gap was evident across multiple dimensions: (i) taxonomic diversity loss, evidenced by consistent reductions in species richness across compartments; (ii) phylogenetic diversity erosion, reflecting the loss of evolutionarily distinct microbial lineages; (iii) functional capacity reduction, with 89% of predicted metabolic functions showing significant alterations with domestication; and, (iv) network fragmentation, demonstrated by the dramatic reduction from 80 to 49 functional connections between teosinte and maize microbiomes. This framework provides a quantitative approach for assessing the effects of domestication on crop microbiomes and could guide restoration efforts in modern breeding programs.

### Functional Erosion of Compartment-Specific Microbiomes

Functional partitioning of the microbiome was evident across plant compartments and the degree of niche specialization diverged markedly between hosts [136]. In teosinte, the mucilage and root-associated communities were dominated by keystone taxa with well-documented plant growth-promoting traits, such as *Azospirillum*, *Bradyrhizobium*, *Herbaspirillum*, and *Pseudoduganella*. These taxa formed tightly connected modules associated with nitrogen fixation, siderophore production (0.992 ± 0.048 in teosinte mucilage), and auxin biosynthesis, supported by a highly modular network structure with a superior clustering coefficient (0.574) and modularity (0.636) [90–92]. In contrast, maize mucilage communities were depleted in these taxa, and were overrepresented by stress-tolerant, fast-growing generalists, such as *Pseudomonas fluorescens*, *Stenotrophomonas maltophilia*, and *Bacillus subtilis*, which may compensate for functional losses with metabolic plasticity [93,94]. Such ecological shifts suggest a transition from a functionally specialized, mutualistic microbiome to a generalist-dominated, opportunistic assemblage, with potential consequences for resilience under low-input or stressful environments [95,96].

Interestingly, *Rhizobium leguminosarum* and *Burkholderia cepacia*, abundant in the rhizosphere of teosinte, were nearly absent in maize, despite their demonstrated contributions to phosphate solubilization and pathogen suppression [97]. Their disappearance, along with the observed reduction in pathogen suppression functions from 0.937 ± 0.065 in teosinte rhizosphere to moderate levels in maize, points to a functional erosion in maize root systems, consistent with earlier observations in soybean and rice [98,99].

### Network Rewiring and Loss of Microbial Integration

The restructuring of microbial functional networks across compartments revealed a stark contrast between teosinte and maize. Microbiomes associated with teosinte exhibited significantly higher network density (0.19 ± 0.01 vs. 0.13 ± 0.01 in maize, P < 0.001), clustering coefficients (0.62 ± 0.03 vs. 0.25 ± 0.02, P < 0.001), and transitivity (0.49 ± 0.02 vs. 0.29 ± 0.02, P < 0.001), particularly in belowground compartments. These features suggest ecologically cohesive communities with strong functional integration. In teosinte mucilage, nitrogen fixation modules formed dense hubs linked to auxin production, phosphate solubilization, and siderophore biosynthesis, involving tightly coupled interactions among *Azospirillum*, *Bradyrhizobium*, *Herbaspirillum*, and *Gluconacetobacter*. These interactions are indicative of highly adapted diazotroph-auxotroph guilds shaped by evolutionary pressures under nutrient-limited environments [100,101].

In contrast, maize microbiomes displayed network fragmentation, with 49 functional connections compared to 80 in teosinte, and a 2.5-fold reduction in clustering coefficient, indicating severely diminished modularity, especially in mucilage and seed compartments. Although an apparent increase in edge density was detected in some networks, this connectivity lacked compartmental structure, suggesting a shift from functionally cooperative modules to generalized, redundant associations [137]. The microbial scaffolds present in teosinte were replaced by more stress-tolerant taxa in maize, such as *Pseudomonas fluorescens*, *Stenotrophomonas maltophilia*, and *Bacillus subtilis*, whose ecological roles may be broader but less specialized [102,103]. This fragmentation implies that Balsas teosinte domestication led to a loosening of microbial interdependence in maize, replacing synergistic consortia with more competitive or self-sufficient assemblages [104].

Notably, taxa such as *Sphingomonas*, *Pantoea*, and *Microbacterium* displayed contrasting roles between hosts. In teosinte, they were embedded within networks linked to phytohormone synthesis and redox metabolism, whereas in maize they appeared disconnected or repositioned within stress-response nodes [105,106]. This suggests a case of niche compression and role reallocation, where a few resilient taxa dominate a wider functional space but with reduced coordination across pathways. The ecological cost of this simplification is likely high as it undermines both redundancy and functional synergy, hallmarks of resilient microbial ecosystems [107–109].

The correlation structure across compartments also revealed that mutualistic functions like nitrogen fixation, plant growth promotion, and phosphorus mobilization were positively associated in teosinte, whereas in maize, these functions became uncoupled or negatively correlated [110]. This decoupling implies a shift in microbial ecology from finely tuned cooperation to functional dispersion, potentially driven by reduced complexity of host-derived substrates or loss of co-evolved microbial traits [111,112].

### The Microbiome as a Casualty of Human Selection for Observable Traits

The taxonomic and functional analysis of microbial communities across compartments highlights that domestication yielded a broad contraction of microbial diversity, particularly in seeds and mucilage [113]. Our recruitment efficiency analysis demonstrates that teosinte maintained significantly superior capacity (1.22 ± 0.18) for assembling diverse microbial communities compared to maize (0.95 ± 0.05), reflecting compromised co-evolutionary relationships disrupted with domestication. The seed microbiome in teosinte maintained a diverse and functionally rich community including *Pseudoduganella*, *Pluralibacter*, and *Variovorax*, taxa associated with quorum sensing interference, redox balancing, and auxin signaling. These organisms not only influence microbial colonization dynamics but also regulate plant immune responses and hormonal homeostasis [85,105].

In maize, those taxa were absent or drastically reduced and seemingly replaced by generalist bacteria, such as *Bacillus* and *Pseudomonas*, which likely offer limited functional breadth. This taxonomic turnover was accompanied by a reduction in mutualistic functions and an enrichment in stress-related pathways [114], which may lead to a microbial system adapted not for diversity and cooperation but for persistence under perturbation and simplified plant-microbe interaction networks. Such restructuring suggests that microbial inheritance was neglected during artificial selection, and with it, the early colonization of beneficial microbes critical for seedling performance and development may have been compromised [115,116].

Functional simplification was especially evident in traits such as siderophore production, auxin biosynthesis, and antifungal metabolite pathways in maize, which were significantly reduced in frequency and centrality [117,118]. While enrichment of stress tolerance and sporulation traits may partly compensate for functional losses, these traits do not fulfill the same ecological functions, indicating that the restructured maize microbiome lacks functional equivalence with that of its wild ancestor [103,119,120].

These findings align with the “microbial domestication cost” hypothesis, which posits that human selection for observable phenotypes such as yield, growth, and grain qualities, inadvertently eliminated less evident, but essential, microbial traits [121,122]. Incidentally, blockage of vertical transmission routes in commercial seeds may explain why modern, commercial maize varieties (hybrids) often require inoculants to perform optimally in enriched environments, while landraces and wild species are inherently more self-sufficient in poor environments through microbe-mediated buffering [123,124]. The systematic nature of these changes—affecting 89% of microbial functions across all plant compartments—suggests that restoring beneficial plant-microbe interactions in modern crops may require comprehensive approaches that consider whole plant-associated microbial networks rather than focusing on individual beneficial microbial strains.

## Conclusion

Our findings indicate that domestication of Balsas teosinte yielded a fundamentally restructured microbiome in maize, likely by disrupting ecological and evolutionary associations with key microbial taxa evident in Balsas teosinte, maize’s wild ancestor. Despite growing under identical conditions, maize and teosinte harbored microbiomes that differed not only in taxonomic composition but also in network complexity, functional specialization, and vertical inheritance. These changes are likely part of a broader pattern of microbial erosion driven by artificial selection, where mutualistic, co-adapted communities have been replaced by communities of generalists with reduced ecological integration. Significantly, domestication-associated microbial erosion—if it has occurred—parallels both domestication [138, 139] and post-domestication [140] genetic erosion in maize driven by artificial and natural selection, further implicating plant genotypic control over its microbiome. The loss of core diazotrophs, hormone modulators, and seed-transmitted taxa suggests that the microbiome is an overlooked casualty of artificial selection, and by extension, systematic breeding as well. Reintegrating such lost functions through targeted microbial restoration and microbiome-conscious breeding offers a path forward to enhance crop resilience, reduce input dependency, and reconnect modern agriculture with its microbial roots.

## 6. Author statements

### 6.1 Author contributions

Conceptualization: J.S.B., E.D.V.C., S.A.B., Data curation: E.D.V.C, S.S.P., S.A.B., Formal analysis: E.D.V.C., S.S.P., Funding acquisition: J.S.B., S.A.B., Investigation: E.D.V.C., S.S.P., J.S.B., S.A.B. Methodology: S.A.B., J.S.B., Project administration: S.A.B., J.S.B., Resources: S.A.B., J.S.B., Software: E.D.V.C., S.S.P., Supervision: S.A.B., J.S.B., Validation: S.A.B., J.S.B., Visualization: E.D.V.C., Writing – original draft: E.D.V.C., S.S.P., Writing – review & editing: E.D.V.C., S.S.P., J.S.B., S.A.B.

### 6.4 Ethical approval

*NA*

### 6.5 Consent for publication

*Mandatory for all journals where personal details of an individual that may lead to their identification has been included in the article. Details include direct identifiers such as names, images and videos; or indirect identifiers that when used together may reveal the individual’s identity (e.g., gender, age, location of treatment, rare disease, socioeconomic data)*.

## Acknowledgements

We are grateful to Messrs. Teodoro Santos Palacios and Alfonso Zarate Zarate for their assistance during field collections. We also thank undergraduate students Arturo España and Amrita Gabu, as well as graduate student Nihad Kerrour, for providing technical support in the laboratory.

