## Supplementary figures and images for "Microbiome differentiation between micro-sympatric maize and teosinte reveals domestication-driven functional erosion of the microbiome across plant compartments"

### Supplementary Figures.pdf

#### Supplementary material

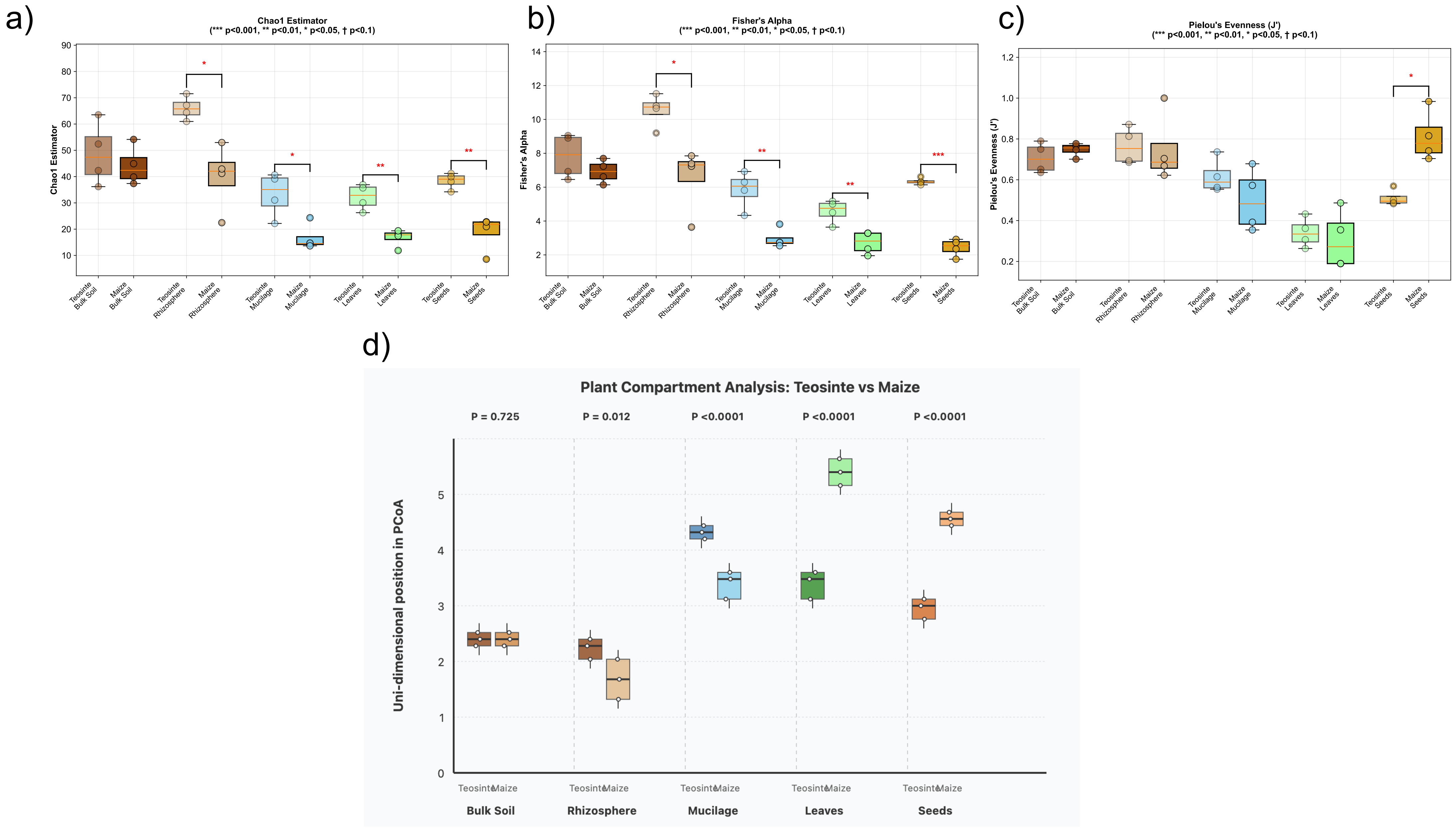
